# Repeated duplications and losses shaped SMC complex evolution from archaeal ancestors to modern eukaryotes

**DOI:** 10.1101/2024.01.07.573240

**Authors:** Jolien J.E. van Hooff, Maximilian W.D. Raas, Eelco C. Tromer, Laura Eme

**Affiliations:** Unité d’Ecologie Systématique et Evolution, CNRS, Université Paris-Saclay, AgroParisTech Gif-sur-Yvette, France; Laboratory of Microbiology, Wageningen University and Research, The Netherlands; Theoretical Biology and Bioinformatics, Faculty of Science, Utrecht University, The Netherlands; Hubrecht Institute, Royal Netherlands Academy of Arts and Sciences (KNAW), The Netherlands; Oncode Institute, Utrecht, The Netherlands; Cell Biochemistry, Groningen Biomolecular Sciences & Biotechnology Institute, Faculty of Science and Engineering, University of Groningen, The Netherlands

**Keywords:** chromosome organization, genome organization, Structural Maintenance of Chromosomes, LECA, condensin, cohesin, SMC5/6, evolutionary cell biology, eukaryogenesis, chromatin, chromosome organization, genome organization, Structural Maintenance of Chromosomes, LECA, condensin, cohesin, SMC5/6, evolutionary cell biology, eukaryogenesis, chromatin

## Abstract

Chromosome organization ensures accurate DNA replication, segregation, gene regulation and DNA damage repair. Across the tree of life, protein assemblages termed Structural Maintenance of Chromosomes (SMC) complexes determine chromosome organization. Eukaryotes usually have four SMC complex types (condensin I, condensin II, cohesin, and SMC5/6), whereas prokaryotes mostly have one. The expanded set is probably needed to accommodate the considerably larger eukaryotic genomes. Despite their essential functions, SMC complexes exhibit notable variation among model organisms, suggesting underexplored diversity across eukaryotes. Here, we provide a thorough reconstruction of the evolution of SMC complex subunits and accessory proteins across eukaryotes. We show that the last eukaryotic common ancestor (LECA) had all four complete SMC complexes, supporting the notion that LECA was already a sophisticated cell. At later timepoints, condensin II was lost at least thirty times, rendering it one of the most frequently lost eukaryotic cellular machineries. We identify multiple components (e.g., Sororin, Securin, Nse5, and Nse6) as much more ancient and widespread than previously appreciated. Finally, we traced the prokaryotic origins of these complexes and propose an ancient SMC complex was already duplicated in the ancestor of TACK and Asgard archaea, suggesting a sophisticated chromosome organization in the archaeal ancestor of eukaryotes. The eukaryotic SMC complex inventory was further expanded through gene duplications, highlighting the importance of these events in the emergence of eukaryotic complexity. Altogether, our results address questions and raise new ones about how SMC complex evolution affected the genome organizations of ancestral and contemporary organisms.

## Introduction

The organization of large eukaryotic nuclear genomes is a complex and dynamic process^1^. At the lowest level, DNA is wrapped around a histone octamer, forming a nucleosome (Figure 1). The DNA then forms topologically associating domains (TADs), wherein DNA sequences interact more frequently with each other than with other regions, with consequences for the regulation of gene expression^1^. At a higher organizational level, chromosomes consist of two main types of spatially distinct compartments, termed “A” (transcriptionally active euchromatin) and “B” (silenced heterochromatin) compartments. At the highest structural level, chromosomes typically occupy their own region in the nucleus. To ensure efficient and faithful DNA replication, maintaining genome stability and minimizing DNA damage, and to allow intricate regulation of gene expression, these organizational levels need to be carefully coordinated.

**Figure 1.**
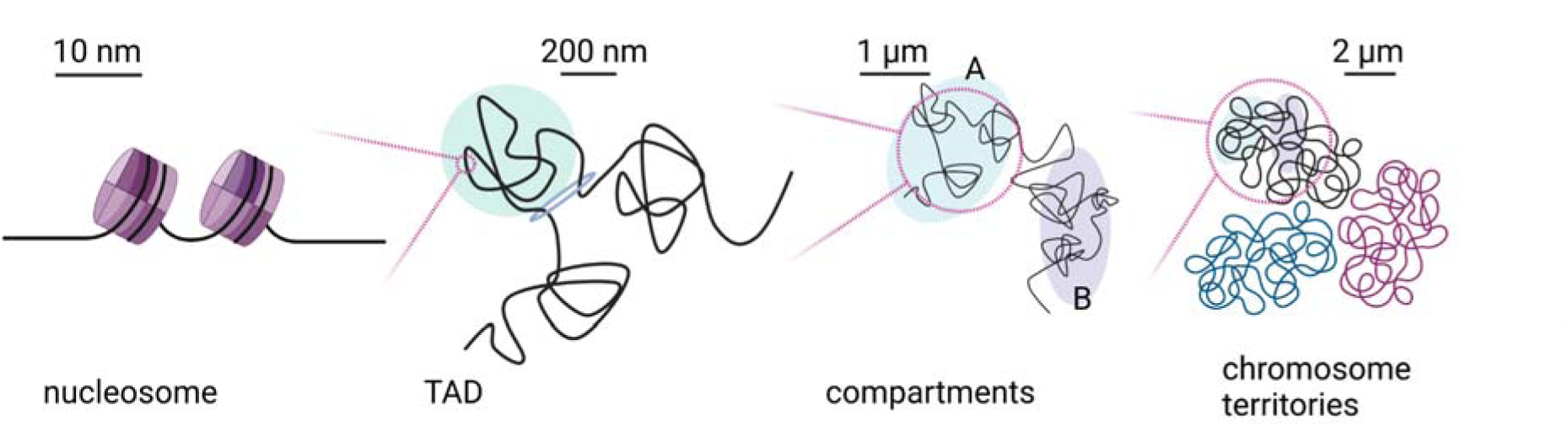
Schematic representation of hierarchical nuclear genome organization in eukaryotes. The typical structural unit of eukaryotic genome organization is the nucleosome, in which 147 base pair are surrounding a histone octamer. Longer stretches of DNA form TADs, regions exhibiting higher interaction frequencies within them compared to adjacent regions. At a higher level, chromosomes are partitioned into distinct A and B compartments. A compartments are enriched with transcriptionally active euchromatin, while B compartments contain repressed heterochromatin. Finally, individual chromosomes are confined to specific subspaces within the nucleus, known as chromosome territories. Created with BioRender.com.

SMC (Structural Maintenance of Chromosomes) complexes have versatile and crucial roles in regulating these different levels. SMC complexes interact with DNA by hydrolyzing ATP. This drives a conformational change that allows it to capture DNA and then move it through the complex. With that, it builds and progressively enlarges DNA loops, a process referred to as loop extrusion^2^. This ability underlies many of the functions of SMC complexes, facilitating the spatial and functional organization of the genome.

SMC complexes have been identified in all three domains of life^3^, and have been shown to function in chromosome condensation, genetic recombination and DNA replication and repair. In addition, in eukaryotes, they are responsible for sister-chromatid cohesion^4^. Eukaryotes typically possess four SMC complexes: condensin I, condensin II, cohesin, and SMC5/6, whereas most prokaryotes have just one (Figure 2B). All SMC complexes share a common basic structure, consisting of two SMC proteins that dimerize via their hinge domains to form a composite ATPase (Figure 2A). A long and flexible kleisin protein binds asymmetrically to one SMC protein’s neck, and to the other’s cap, resulting in a tripartite ring that is supplemented with different proteins, depending on the complex type. Eukaryotic condensin I and II, and cohesin contain hawk proteins, while prokaryotic SMC complexes and the eukaryotic SMC5/6 complex contain kite proteins. Hawks and kites do not appear to be homologous^5^. Eukaryotic SMC complexes also incorporate accessory, sometimes transient subunits, which regulate their activity and localization.

**Figure 2.**
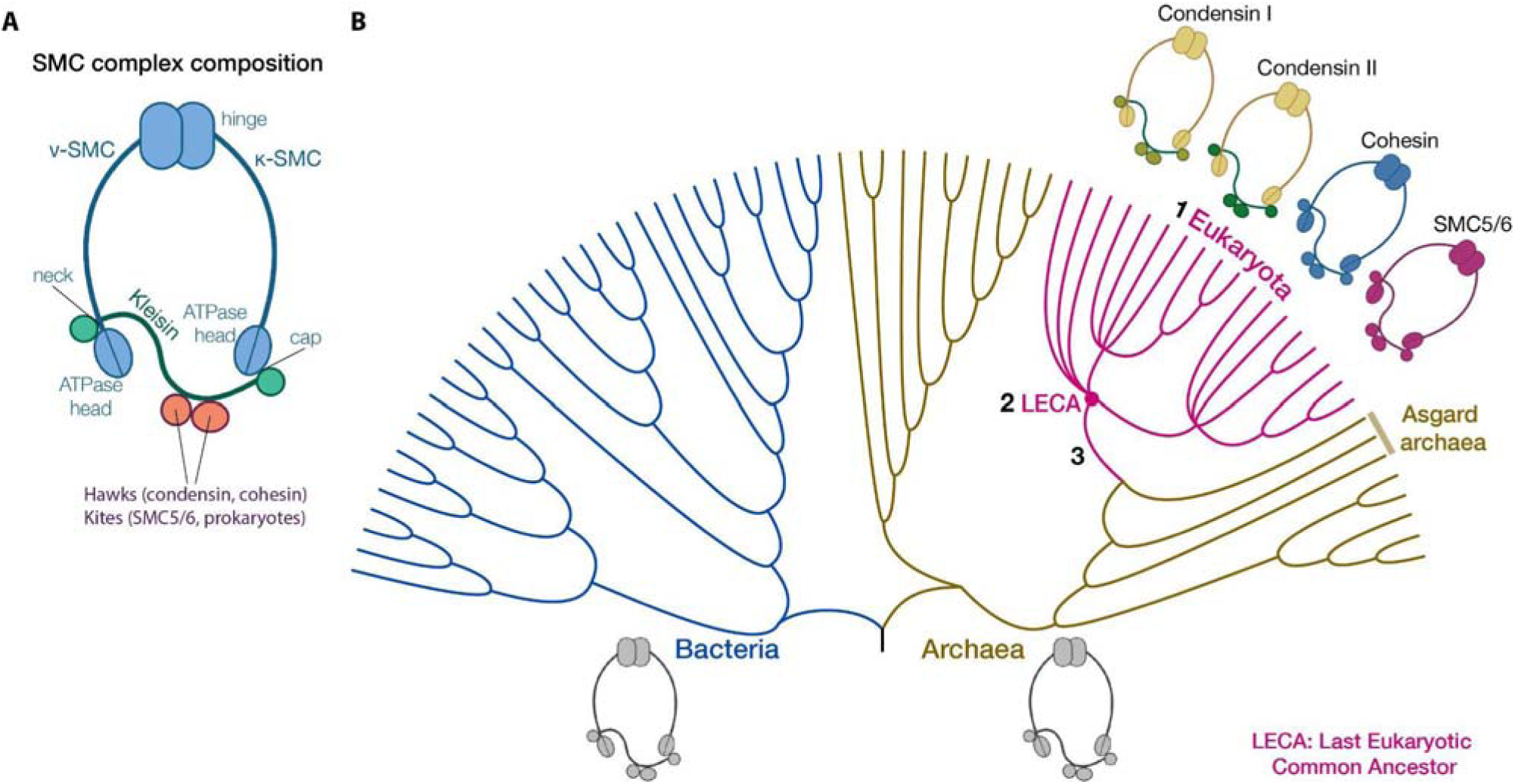
SMC complex composition and evolution. **A**. General composition and architecture of SMC complexes across the tree of life. The SMC proteins form dimers via their hinge domains (top) and their ATPase heads are linked to the kleisin protein (bottom). Between the head and hinge domains, they feature a backfolded coiled-coil region. The kleisin attaches to one SMC protein through its N-terminus, binding the SMC protein at its neck (the _ν_-SMC), and to the other SMC protein via its C-terminus, binding it at the SMC protein’s cap (the _κ_-SMC). Note that in prokaryotic SMC complexes, the _ν_-SMC and _κ_-SMC are the same protein, forming a homodimer. In eukaryotic SMC complexes, these are different proteins forming a heterodimer. In addition to the SMC and kleisin proteins, SMC complexes generally include two other core subunits, either hawk proteins (the eukaryotic condensin and cohesin complexes) or kite proteins (prokaryotic SMC complexes and the eukaryotic SMC5/6 complex). Eukaryotic SMC complexes often have accessory members and, or more transient interactors (Figure 3). **B**. Illustration of the tree of life with this study’s research questions. Eukaryotes typically have four SMC complexes: condensin I, condensin II, cohesin and SMC5/6. Research questions: (1) What is the diversity of SMC complexes among eukaryotes? (2) What were the compositions of the SMC complexes in the Last Eukaryotic Common Ancestor (LECA)? (3) How did eukaryotic SMC complexes and their subunits originate before LECA? Created with BioRender.com.

Each complex has distinct roles, but there are areas of overlap, and several functions remain unclear. Condensin I and II complexes, which condense chromosomes during mitosis^6^, are composed of identical SMC proteins (SMC2 and SMC4) but differ in their kleisin (CAPH versus CAPH2) and hawk proteins (CAPG and CAPD2 versus CAPG2 and CAPD3, respectively) (Figure 2A). The impact of condensin II extends to interphase, as it induces the formation of chromosome territories^7^. This appears consistent across studied species, as those lacking condensin II do not have territories. Instead, they exhibit a distinct nuclear arrangement known as a ‘Rabl’-like architecture, characterized by the clustering of centromeres and telomeres respectively, and often their attachment to the nuclear envelope^8^.

The primary role of cohesin is in maintaining sister chromatids in cohesion from DNA replication until anaphase, ensuring chromosomes are properly aligned and attached to the mitotic spindle for safe segregation. Furthermore, at least in vertebrates, cohesin organizes DNA into loops, which in turn form TADs. Cohesin’s composition seems more intricate than that of condensin, characterized by a diverse array of core and peripheral subunits (Figure 3).

**Figure 3.**
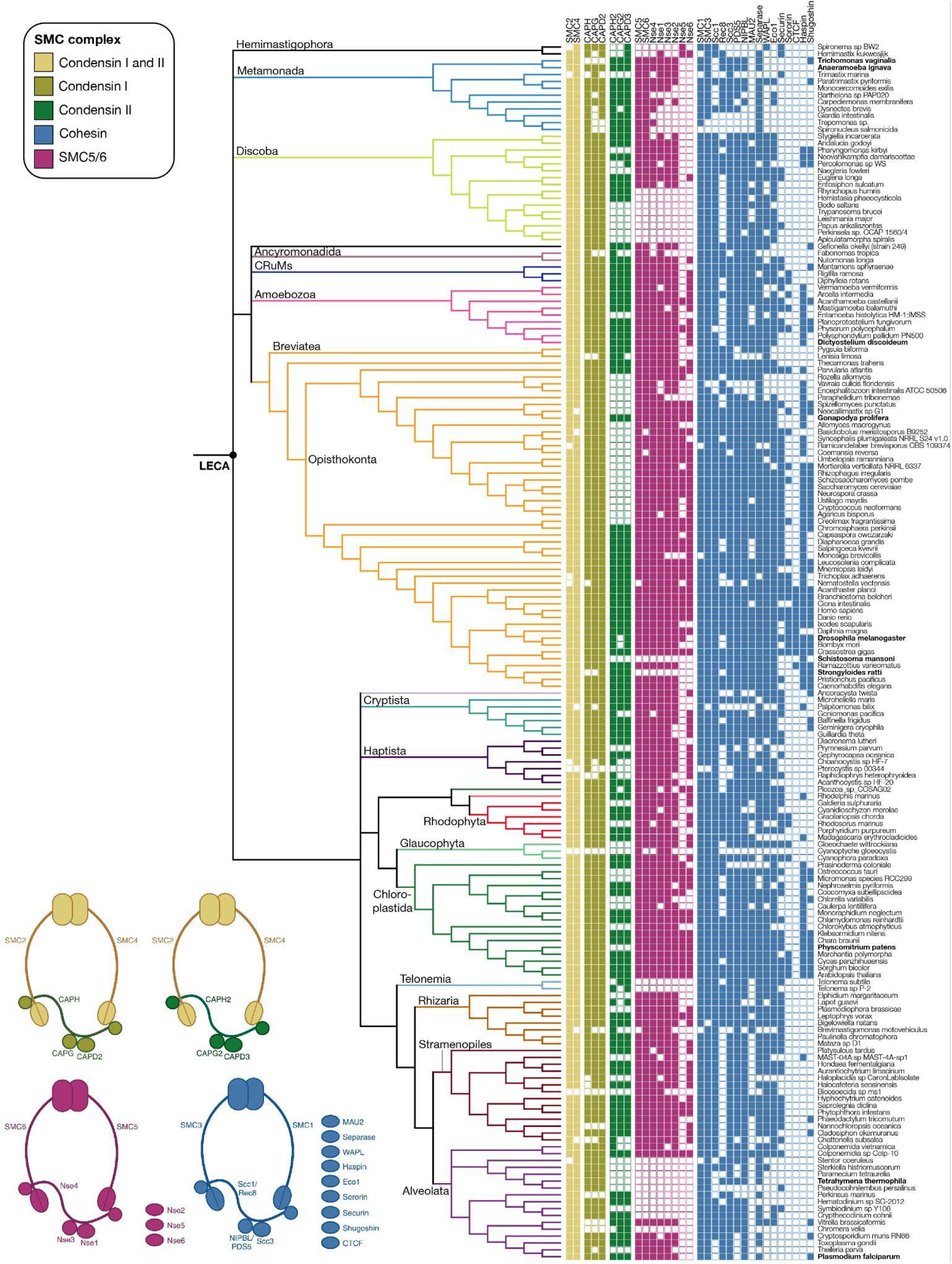
Distributions of SMC complexes’ subunits across the eukaryotic tree of life. Species referred to in the text are displayed in bold. Bottom: illustrations of the eukaryotic SMC complexes with their subunits. **Condensin I and II** have the SMC proteins SMC2 and SMC4, kleisin protein CAPH (condensin I) or CAPH2 (condensin II) and hawk proteins CAPG and CAPD2 (condensin I) or CAPG2 and CAPD3 (condensin II). The **SMC5/6** complex has the SMC proteins SMC5 and SMC6, kleisin protein Nse4 and kite proteins Nse1 and Nse3. It also contains Nse2, Nse5 and Nse6. **Cohesin** has the SMC proteins SMC1 and SMC3, kleisin proteins Scc1 (mitosis) or Rec8 (meiosis), hawk proteins Scc3, PDS5 and/or NIPBL. Other (transient) subunits and interactors include MAU2, WAPL, Eco1, Securin, Sororin, Haspin, Shugoshin, Separase and CTCF. Created with BioRender.com.

Finally, the SMC5/6 complex stands out as the most cryptic eukaryotic SMC complex. In addition to the core elements (Figures 2A, 3), it contains a SUMO ligase (Nse2) and a helix-rich subcomplex (formed by Nse5 and Nse6)^9^. The SMC5/6 complex is implicated in a variety of cellular processes, including genome replication, DNA damage repair, and defense against viral DNA^9^.

The absence of some SMC complexes has been reported in distantly related eukaryotic lineages^10–12^, but their precise distribution and composition, evolutionary history during the diversification of eukaryotes, and impact on genome organization in non-model organisms remain unknown. In addition, the evolutionary events leading to the expansion of the SMC complex repertoire in eukaryotes remains to be precisely determined. Yoshinaga & Inagaki suggested that two SMC complexes were already present in the Asgard archaeal ancestor of eukaryotes^13–15^, which later, in early eukaryotic evolution, gave rise to condensin plus cohesin, and to SMC5/6, respectively^10^. However, the sequence of evolutionary events leading to the emergence of the four eukaryotic SMC complexes and their distinct subunits remains unresolved. Specifically, the evolution of non-SMC protein components, including the kleisin, kite and hawk families, has not yet been studied. As a result, it is not known yet whether the archaeal ancestor of eukaryotes indeed possessed two distinct, complete SMC complexes.

Here, we systematically investigate the distribution and evolution of eukaryotic SMC complexes, both before and after the last eukaryotic common ancestor (LECA). We reconstruct the SMC repertoire of LECA, uncover how these complexes diversified post-LECA, and propose a scenario for the origin of all four SMC complexes pre-LECA. Our analysis is based on state-of-the-art homology detection, including structural comparisons, and molecular phylogenetics, applied to a diverse dataset of 177 eukaryotic and 4,101 prokaryotic proteomes, including 28 Asgard archaea—the closest extant prokaryotic relatives of eukaryotes^15^.

Our findings shed new light on the evolution of these hallmark eukaryotic protein complexes, provide insights into genome organization in non-model eukaryotes, and offer a valuable framework for in-depth functional investigation of individual proteins through their sequence evolution.

## Results

### Condensin II lost recurrently during eukaryotic evolution

We first searched for orthologs of condensin I and II subunits across a diverse range of eukaryotes (Methods). We confirmed that SMC2 and SMC4 are widely conserved, suggesting that the vast majority of species possess at least one condensin complex (Figure 3, Supplementary Table 1). Our analysis revealed that the subunits specific to each complex, i.e. CAPH, CAPG, and CAPD2 (condensin I), and CAPH2, GAPG2, and CAPD3 (condensin II), are generally coherently present or absent, suggesting their co-evolution across eukaryotic diversity.

The absence of complex-specific subunits in various eukaryotic species suggest multiple independent losses of condensin I and II (at least six and thirty, respectively). To avoid false negatives owing to extensive sequence divergence or incomplete data, we only considered a condensin complex absent from a given lineage if all three complex-specific proteins were undetected. Accordingly, condensin II subunits are among the most frequently lost proteins in eukaryotic evolution^16–19^, matching several kinetochore proteins^20^ and a cilium-associated small GTPase^21^. The absence of condensin II has been reported to correlate with a Rabl-like architecture^12^ (although exceptions exist, see Discussion) and the remarkably high frequency of condensin II losses suggests that transitions from chromosome territories to Rabl-like interphase nuclear architectures have been common during eukaryotic evolution. Moreover, it suggests that in contrast to condensin I, condensin II is often dispensable for condensation of chromosomes during mitosis.

Conversely, we report for the first time that certain lineages exclusively possess condensin II, such as the human pathogen *Trichomonas vaginalis*. While we identified condensin I subunits in its relative *Anaeramoeba ignava*^22^, thorough sequence profile- and structure-based homology searches did not yield any candidate condensin I subunits in this species (Figure 3). Importantly, few relatives of *T. vaginalis* have been sequenced, impeding a robust comparative approach and a more precise pinpointing of the loss of condensin I.

Additionally, whereas condensin II was thought to be entirely absent in fungi^23^, we detected it in the fungus *Gonapodya prolifera*, and in its sister species *Gonapodya sp.* JEL0774, confirming that the presence of condensin II was not due to contamination. Although a possibility, our protein phylogenies do not unequivocally demonstrate that these species acquired condensin II via horizontal gene transfer (HGT; Supplementary Data), leaving the exact evolutionary history of condensin II in Fungi unclear. Regardless, the presence of condensin II in *Gonapodya* raises the question of the existence of chromosome territories in this genus.

Finally, we confirm that the ciliate *Tetrahymena thermophila* expanded its condensin I repertoire in the absence of condensin II^24,25^. Although the phylogenies of CAPH, CAPG and CAPD2 proteins are not well-resolved (Supplementary Data), they suggest that these proteins may have duplicated in parallel across different ciliate lineages^25^. Upon functional differentiation, these expanded repertoires likely enable ciliates to independently orchestrate their distinct somatic and germline genomes.

### LECA harbored a complete SMC5/6 complex, including Nse5 and Nse6

Our analyses strongly suggest that LECA harbored a complete SMC5/6 complex, including the core subunits SMC5, SMC6, kleisin Nse4, and the kites Nse1 and Nse3, as well as the accessory proteins Nse2, Nse5 and Nse6. Our expanded taxon sampling suggests that the SMC5/6 complex was lost at least eight times during eukaryotic evolution (Figure 3, Supplementary Table 1). These losses range from ancient events, such as in the ancestor of ciliates, which may have lived over 1 Ga years ago^26^, to more recent occurrences, including in the metazoan blood fluke *Schistosoma mansoni* and the rat parasite *Strongyloides ratti*.

Interestingly, the primary subunits, SMC5,6 and Nse1,3,4, along with the associated SUMO ligase Nse2, consistently co-occur across species, suggesting that Nse2 is integral to the SMC5/6 complex, functioning as a core component since before LECA, and that the complex’s sumoylation activity is an ancient feature.

Nse5 and Nse6 were previously thought to be taxonomically restricted and pose unique challenges for ortholog detection owing to their rapid sequence divergence. Consequently, direct sequence comparisons did not identify homology between the fission yeast Nse5 and its human counterpart SLF1^27^, which were considered functional analogs, not homologs. However, using sensitive sequence searches, structure prediction and similarity analyses, we found that Nse5 counterparts in diverse model organisms are in fact homologous. The same is true for Nse6 equivalents. Remarkably, we identified a putative Nse5 in *Drosophila melanogaster* (gene Dmel\CG5664), a species previously thought to lack Nse5 and Nse6^28^. Using structural similarity searches, we confirmed that the *D. melanogaster* Nse5 candidate strongly resembles Nse5 proteins in human, mouse and fission yeast, and is present in a range of Diptera species (Methods).

Despite detecting Nse5 and Nse6 across a diverse range of eukaryotes, their distributions remain sparse in species possessing other SMC5/6 subunits. This may reflect the difficulty in identifying such highly divergent orthologs, rather than true absences. This divergence is also exemplified by the additional domains found in Nse5 and Nse6 in various lineages, like ankyrin and BRCT repeats in some Nse5 orthologs^29^. Moreover, Nse5 and Nse6 adopt a similar alpha-solenoid structure (Supplementary Figure 2), suggesting a shared evolutionary origin. Based on their structural similarities, we suggest that they resulted from a single ancestor protein that duplicated before LECA (Supplementary Text).

### Cohesin displays a mosaic of divergent members and partners

The presence of SMC1 and SMC3 in nearly all eukaryotes analyzed suggests a near-ubiquitous distribution of cohesin (Figure 3, Supplementary Table 1). However, the distribution of non-SMC subunits is more sporadic and inconsistent, with different subunits absent in various lineages, obscuring any clear functional and evolutionary modules among these components.

Cohesin includes two kleisins, Scc1 and Rec8, functioning in mitosis and meiosis, respectively^30^. The wide but patchy distributions of these paralogs (Figure 3), combined with their phylogeny (Supplementary Data), suggest that Scc1 and Rec8 resulted from a pre-LECA gene duplication, that both were present in LECA, and were then independently lost multiple times. The limited sequence divergence between Scc1 and Rec8 suggests their functions have remained closely related, with the mitosis-meiosis differentiation possibly incomplete in LECA. This in turn could explain the frequent loss of one of the two paralogs, owing to potential functional redundancy.

The cohesin complex also contains three hawk proteins: PDS5, NIPBL, and Scc3. All three are relatively sparsely distributed, suggesting that they are also partially functionally redundant, in line with the evidence that PDS5 and NIPBL exclude each other from binding to the kleisin^31,32^. Yet, various lineages have multiple paralogs of individual hawks, including humans, which have multiple copies of Scc3 (SA1/2/3) and PDS5 (PDS5A/B). However, these paralogs were derived from recent gene duplications, and we infer that LECA possessed a single copy of each of PDS5, NIPBL and Scc3. Cohesin function is modulated by several regulators, including Securin and Sororin, both crucial for sister chromatid cohesion. Our reevaluation of Securin’s characterized analogs across model organisms confirmed their homology^33^. Similarly, we confirm the homology of candidate Sororin proteins^34^ and their broad distribution. Both of these largely disordered regulators were detected primarily through the presence of short linear motifs, which are not always unique, raising the possibility of convergent evolution in some cases (Supplementary Text). Altogether, the broad presence of Securin and Sororin across eukaryotes suggests that these proteins were already regulating cohesin in LECA.

### Eukaryotes inherited an expanded SMC array from archaeal ancestors

#### SMC proteins

To investigate the pre-LECA origins of the eukaryotic SMC complexes, we performed phylogenetic analyses. These confirm that eukaryotic SMC proteins form two clusters^10,37^, one consisting of condensin and cohesin SMC proteins, and the other by SMC5 and SMC6, separated by several clades of prokaryotic homologs (Figure 4A). However, topology tests suggest that the possibility that these two clusters actually form a monophyletic group cannot be rejected (p-AU=0.317). Both condensin/cohesin SMC proteins and SMC5,6 show archaeal homologs as their closest relatives, including Asgard archaea, indicating that eukaryotes inherited them from their archaeal ancestors (Supplementary Table 3).

**Figure 4.**
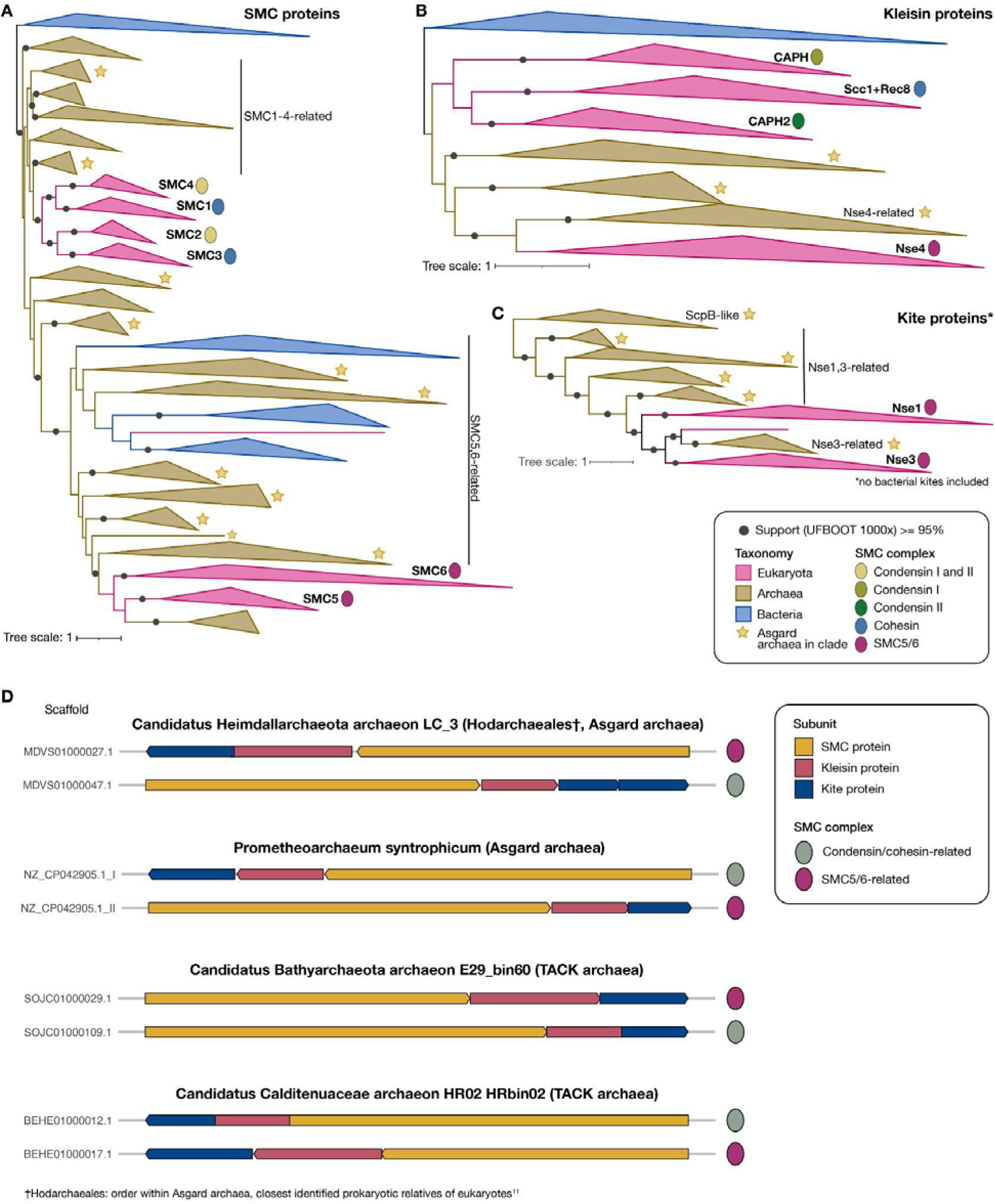
Phylogenetic relationships between eukaryotic and prokaryotic SMC complexes. **A-C.** Phylogenetic trees of SMC (A), kleisin (B) and kite (C) proteins inferred using IQ-TREE^35^ under the LG+C60+G mixture model. Statistical support at branches was estimated from 1000 ultrafast bootstrap replicates^36^. The uncollapsed, unfiltered trees can be found on https://itol.embl.de/tree/13722425212266531697468040 (SMC), https://itol.embl.de/tree/137224252126461697465203 (kleisin), and https://itol.embl.de/tree/13722425212198711697525255 (kite). Scale bars indicate estimated average numbers of substitution per site. **D**. Genomic localizations of SMC complex operons, with genes encoding SMC proteins, kleisins and kites in a selection of archaea having at least two paralogs of each.

Several archaea, particularly TACK and Asgard archaea, have two SMC proteins (Supplementary Table 3), likely resulting from a gene duplication event in an early archaeal ancestor. Our analyses confirm that after eukaryotes diverged from their archaeal ancestors, the ancestral condensin/cohesin SMC protein underwent an additional duplication event, resulting in the ancestors of SMC1,4 and SMC2,3. Similarly, the ancestor of SMC5,6 duplicated to give rise to SMC5 and SMC6^10^. In both cases, the duplication resulted in a distinct κ-SMC (Figure 2A, SMC1,4, and 5) and ν-SMC (SMC2,3, and 6), while in prokaryotes the interacting partners are the same protein. Thus, these duplications and subfunctionalizations within the SMC protein complexes occurred in parallel, yielding SMC protein heterodimers in eukaryotic SMC complexes rather than the homodimers seen in prokaryotes. After this heterodimer-forming duplication, the condensin/cohesin SMC proteins duplicated once more, giving rise to the condensin and cohesin SMC heterodimers.

#### Kleisin proteins

In contrast to previous work^10^, our phylogeny suggests that the kleisins from condensin/cohesin and from SMC5/6 do not form a monophyletic group (Figure 4B). Their monophyly was rejected by the Approximately Unbiased (AU) topology test (p-AU=0.002). Interestingly, the SMC5/6 kleisin, Nse4, branches with strong support together with a clade containing kleisins from TACK and Asgard archaea, suggesting an Asgard archaeal provenance of Nse4, in line with the Asgard archaeal ancestry of eukaryotes^14,15^. By contrast, while we observe other clades containing a wide representation of archaea, they do not form a monophyletic group with condensin/cohesin kleisins.

We hypothesize that these non-Nse4-related archaeal clades, which contain the same organismal diversity as the Nse4-related archaeal clade (i.e., TACK and Asgard archaea), represent the true sister group of the condensin/cohesin kleisins. This is supported by inspection of the genomic context of this group of archaeal kleisins, which shows them encoded next to condensin/cohesin-related SMC proteins (Supplementary Text, Figure 4D). The duplication of kleisins in an archaeal ancestor would be coherent with the duplication scenario for the SMC proteins outlined above. Within the condensin/cohesin kleisins clade, it is not clear which kleisins are most closely related to one another, since these branches are poorly supported. Given the putative close relationship of condensin I and II, which share SMC proteins, we hypothesize that the condensin kleisins CAPH and CAPH2 are more closely related to each other than to the cohesin kleisins; a hypothesis also hinted at by a more restricted phylogenetic analysis (Methods, Supplementary Figure 1).

In summary, eukaryotic SMC proteins and kleisins appear to have originated from an ancient archaeal gene duplication, probably in the ancestor of TACK and Asgard archaea, which was followed by further duplications specific to eukaryotic evolution.

#### Kite proteins

Our identification of prokaryotic kite homologs of Nse1 and Nse3 (SMC5/6 complex) suggests that at the sequence level, kites fall into three distinct archetypes: Nse1/3-like (archaeal kites only), ScpB-like (both archaeal and bacterial kites, such as ScpB in *Bacillus subtilis*), and MukE-like (other bacterial kites, like MukE in *Escherichia coli*). Despite their marked sequence divergence, these archetypes share similar structural features^38^. Our kite phylogeny mirrors that inferred from SMC and kleisin proteins, where various archaea, in particular TACK and Asgard archaea, possess two kites, namely one Nse1/3-like kite, and one ScpB-like kite (Figure 4C). Thus, the kite protein likely duplicated in the ancestor of TACK and Asgard archaea.

Our analyses of SMC protein, kleisin, and kite paralog distributions indicate that many archaea harbor two complete SMC complexes. Examination of the genomic localization of these genes in several representative species shows that the paralogs are organized into distinct operons, each encoding all three subunits (Figure 4D). Based on our phylogenetic inferences, we identify these as condensin/cohesin-related and SMC5/6-related complexes. Although the majority of species possessing both complexes belong to the TACK and Asgard archaeal lineages, a few euryarchaeal species also carry them (Supplementary Table 3), potentially owing to rare HGT events. Given their overall distribution, we propose that the duplication of the entire SMC complex operon likely occurred in the common ancestor of TACK and Asgard archaea.

Of note, some Asgard archaea possess three kite paralogs (including Hodarchaeales^15^), with one of them forming an exclusively Asgard archaeal clade clustering with (eukaryotic) Nse3. This suggests that the duplication event leading to Nse1 and Nse3 occurred in an Asgard archaeal ancestor. Moreover, these paralogous Asgard kite genes co-localize in the genome with different SMC and kleisin genes, supporting their involvement in distinct SMC complexes, and hence suggesting an additional duplication of the whole SMC complex (Supplementary Text).

#### Hawk proteins

Hawk proteins are HEAT-repeat proteins that form part of the condensin and cohesin complexes. We employed extensive homology searches to investigate their deep evolutionary origins, yet were not able to detect hawk proteins in prokaryotes. In addition, the degenerate nature of these repeats renders their sequences too divergent to conduct conventional phylogenetic analysis. Thus, to investigate their relationships to each other, and to other HEAT-repeat containing proteins, we adopted a sequence profile network approach^5^, combined with comparisons of AlphaFold2-predicted 3D structures. This leads us for example to conclude that cohesin release factor WAPL is not a HEAT repeat, but an Armadillo repeat-containing protein (Supplementary Text).

Our analysis confirms that hawks are paralogous to one another^5^, following gene duplication events that occurred before LECA. Interestingly, hawk proteins form two groups in the profile and structure analyses, which correspond to the previously described functional groups in the condensin/cohesin complexes^39^: the A-type hawks (CAPG, CAPG2, Scc3), and the B-type hawks (CAPD2, CAPD3, NIPBL and PDS5) (Figure 5A,B, Supplementary Tables 5,6,7). A likely scenario is that, in the ancestor of condensin and cohesin, a single hawk protein first replaced kite(s). This protein first formed a homodimer, and then duplicated, resulting in an A- and B-type-containing heterodimer.

**Figure 5.**
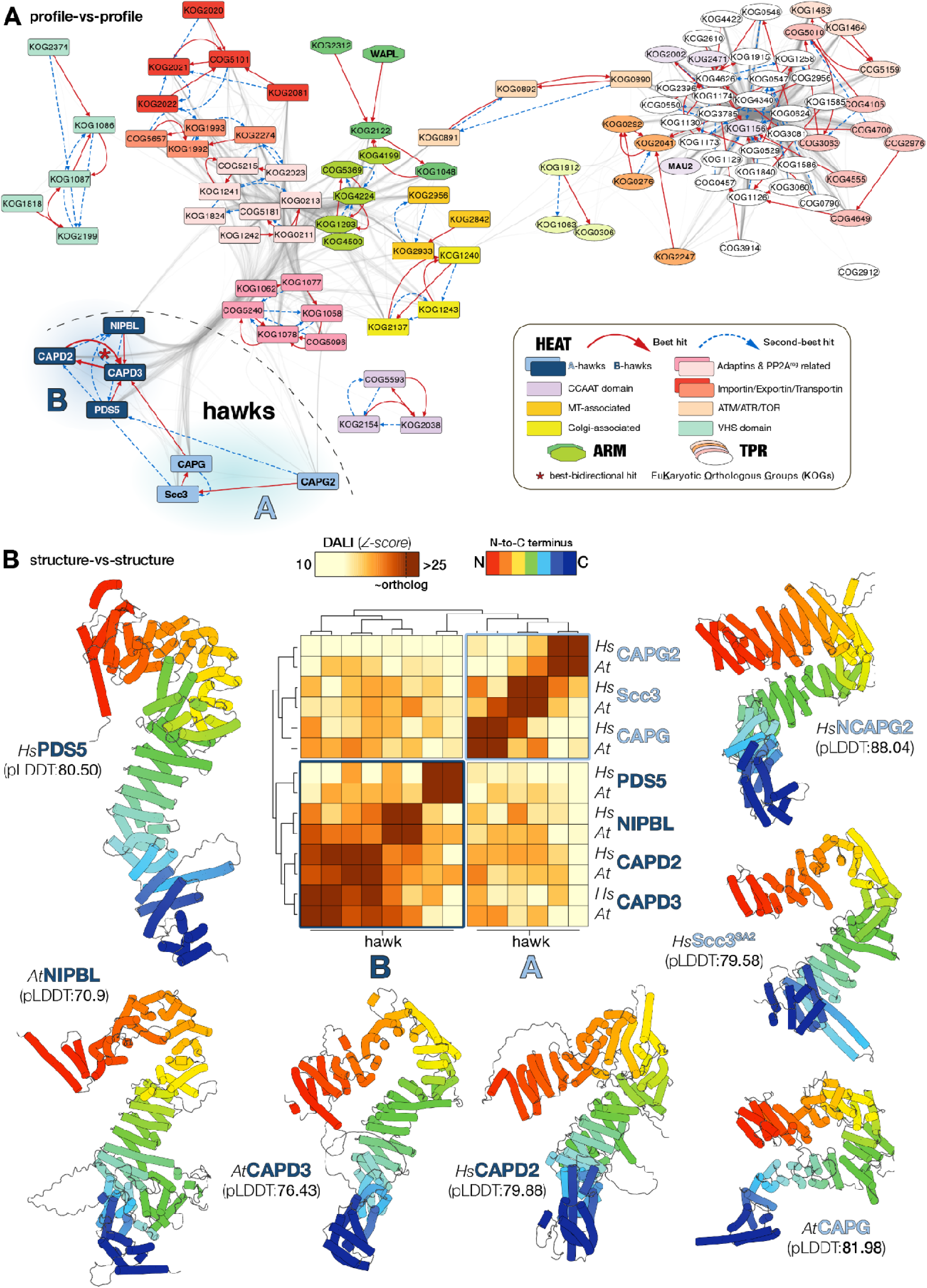
HMM profile network analysis and 3D protein structure comparisons reveal an evolutionary split between A- and B-type hawks. **A**. Graph of a profile-vs-profile-based network analysis. The initial query profiles were those of hawk proteins, WAPL and MAU2, which were searched against the COG/KOG database. All profile hits identified in the initial search were subsequently taken as query, creating a bidirectional graph. Displayed edges are filtered to show only significant hits (E-value <= 1E-3, alignment length >= 100, which corresponds to ∼2 HEAT repeat units). Nodes are colored based on MCL cluster identity (Methods). **B**. Predicted structures of hawk proteins of *Homo sapiens* (*Hs*) and *Arabidopsis thaliana* (*At*) (Methods). Pairwise alignment Z-scores as computed by Dali^40^ are represented in the heatmap, generated with ClustVis^41^.

## Discussion

### An evolutionary scenario for the emergence of eukaryotic SMC complexes

Our phylogenetic analyses of SMC proteins, kleisins and kites suggest an ancient duplication of the entire SMC complex in an archaeal ancestor (Figure 6). We propose that the duplicated SMC proteins formed separate homodimers, rather than a single heterodimeric complex, which is supported by the genomic co-localization of each complex’s candidate subunits in extant archaeal genomes (Figure 4D).

**Figure 6.**
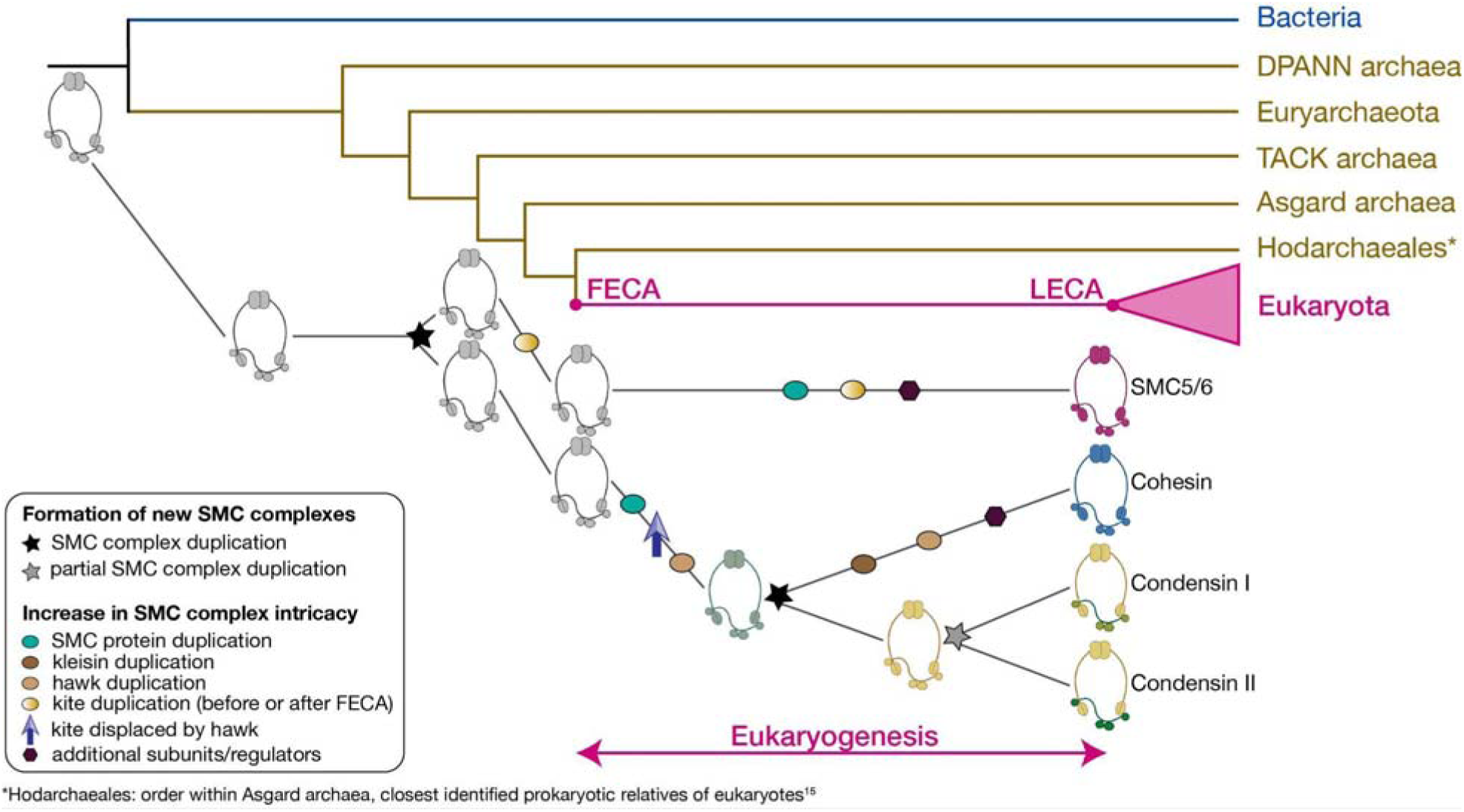
Evolutionary scenario for the origins of the four eukaryotic SMC complexes between the First Eukaryotic Common Ancestor (FECA) and LECA, referred to as ‘eukaryogenesis’. FECA possessed two distinct SMC complexes, a legacy of an early archaeal duplication event of the SMC complex, which we hypothesize to have occurred in the ancestor of TACK and Asgard archaea (most left-sided black star). One of these complexes underwent a duplication of both its SMC and kite proteins, leading to the evolution of the SMC5/6 complex. Note that the timing of the kite duplication is unresolved: it might either have occurred before FECA (it would be shared with Asgard archaea, Figure 4C) or after FECA. The SMC protein duplication transformed the original SMC protein homodimer within the complex into a heterodimer, comprising a _κ_-SMC and a _ν_-SMC. The second complex also underwent a duplication of its SMC protein. However, in this instance, the kite subunits were replaced by hawk proteins. This SMC complex underwent an additional duplication, resulting in the emergence of cohesin and condensin. Subsequently, the condensin complex underwent a partial duplication, specifically affecting its non-SMC proteins, namely kleisin and hawks. Finally, within the ancestral condensin/cohesin, the hawk proteins also duplicated. The timing of the hawk duplications is speculative, as we were not able to conduct a phylogenetic analysis for this family. The composition of SMC complexes is illustrated in Figures 2A,3. Created with BioRender.com.

The SMC complex duplication event likely occurred in the ancestor of TACK and Asgard archaea. We propose that during the origination of eukaryotes (‘eukaryogenesis’), each complex underwent SMC protein duplication and subsequent subfunctionalization, generating heterodimers composed of κ-SMC and ν-SMC subunits (Figure 6). The proposed SMC protein duplication occurred in parallel in both the condensin/cohesin ancestor as well as in the SMC5/6 complex ancestor. Indeed, such homomer-to-heteromer duplications are very common in eukaryotic evolution^42,43^, yet they might not necessarily reflect any selective advantage^44,45^. In the condensin/cohesin ancestor, the ScpB-like kite protein was replaced by a hawk protein. This coincided with kleisin elongation, as condensin and cohesin kleisins are longer than those in the SMC5/6 complex and prokaryotic SMC complexes^5,39^. While phylogenetic analyses of hawk proteins were hampered by their extensive divergence, our network analysis suggests the first hawk formed a homodimer, which likely duplicated and formed a heterodimer of A- and B-type hawks^39^. This complex then duplicated, giving rise to condensin and cohesin. Condensin experienced a partial duplication, affecting only the kleisin and hawks, resulting in condensin I and II. Additionally, the cohesin kleisin and one hawk duplicated. In the SMC5/6 ancestor, the kite protein duplicated, likely before the divergence of eukaryotes from Asgard archaea (Figure 4C).

Beyond these duplication events, the SMC5/6 complex and cohesin recruited additional subunits and regulators (e.g., Nse2, Nse5/6 for SMC5/6; WAPL, Securin, and Sororin for cohesin). Existing members also acquired new domains; for example, Nse1 gained a RING-finger E3 ubiquitin ligase domain^46^, absent in Nse3 and archaeal kites. This eukaryotic innovation may enable SMC5/6 to ubiquitinate proteins^47^, just as the addition of Nse2 enabled it to sumoylate proteins. Combined, our data demonstrate that evolutionary mechanisms including gene duplication, replacement and recruitment endowed LECA with the four SMC complexes found in modern eukaryotes. Furthermore, our analysis of SMC complex evolution highlights the importance of gene duplication in eukaryotic complexity^48,49^, as it involved at least two whole-complex duplications, one partial-complex duplication, and multiple intra-complex duplications.

### Outlook: towards evolutionary cell biology of SMC complexes

The distributions of SMC complexes across eukaryotes raise questions about their functional differences. For example, what roles does condensin I have in species lacking condensin II? Do condensin complexes fulfill any non-mitotic functions, such as forming TADs^50,51^? Do species lacking condensin II exhibit a Rabl-like nuclear architecture rather than a chromosome territory one? Several of the species in our dataset do not conform to the latter correlation. For example, *Plasmodium falciparum* and *Dictyostelium discoideum* possess condensin II and yet have a Rabl-like organization^52,53^, while the moss *Physcomitrium patens* lacks condensin II and forms chromosome territories^54^. Scrutinizing this relationship experimentally would be particularly relevant in Fungi, where we identified condensin II in the chytrid lineage of *Gonapodya*.

Furthermore, exploring lineages where SMC5/6 has been recently lost, like in the nematode *S. ratti*, could enhance our understanding of the dispensability of this complex.

In addition to raising questions about the implications of protein presences and absences, our ortholog sets allow for an in-depth examination of individual protein sequences, which may reveal new sequence motifs mediating functions, for example via protein-protein interactions. As our work comprises a wide range of SMC complex proteins, many different researchers might profit from this possibility. It is important to point out that what constitutes an SMC complex protein is somewhat blurred, with some proteins being only loosely associated or transient interactors, which we included in order to offer a comprehensive overview.

The relationship between nuclear chromosome architecture and condensin II remains to be confirmed across a wider, more diverse range of species. If true, it would pave the way to address questions about the cellular consequences of the frequent losses of the condensin II complex. In particular, transitioning from chromosome territories to a Rabl-like architecture, as a result of loss of condensin II, may lead to decreased chromatin flexibility owing to chromosome tethering to the nuclear envelope^55^, and therefore reduce precise and context-dependent control of gene expression. Yet, the functional role of chromosome territories has been questioned^56^. At the same time, a Rabl-like configuration might allow cells to accelerate their cell cycle, as it requires less reorganization before going into mitosis^57^. Such a configuration could be especially advantageous in life histories involving rapid cell divisions, as seen in certain parasitic or rapidly growing fungi.

Future investigations comparing closely related species or strains that differ in condensin II presence would help dissect whether (simpler) nuclear organization correlates with faster cell cycles—and possibly diminished regulatory plasticity. Such studies could use imaging or Hi-C-based approaches to explore how nuclear architecture dynamics influence gene expression, replication timing, and genomic stability. Ultimately, clarifying the balance between these potential benefits (e.g., faster proliferation) and costs (e.g., reduced chromatin flexibility) could illuminate how chromosome configuration evolves in diverse eukaryotic lineages.

Our findings suggest that diverse archaea, particularly within the TACK and Asgard groups, harbor both condensin/cohesin-related and SMC5/6-related complexes. This hypothesis can be directly tested through experimental investigation of extant archaea to confirm the presence and functionality of these two distinct SMC complexes *in vivo*. Such studies not only promise to validate our predictions but also offer a unique opportunity to elucidate the ancestral functions of condensin/cohesin and SMC5/6. The characterization of the SMC5/6-related complex in archaea is of particular interest, given the ongoing uncertainty regarding its canonical eukaryotic function.

Altogether, the evolutionary trajectories of SMC complexes provide a starting point for studying the genetic underpinnings of eukaryotic genome organization (Figure 1). In particular, they might help to understand how and why eukaryotes differ in their genome organizations, and how their organizational complexity emerged during eukaryogenesis^58^.

## Methods

### Eukaryotic dataset assembly

To determine the presences and absences of eukaryotic SMC complex subunits across the eukaryotic tree of life, and to reconstruct their evolutionary histories, we used a eukaryotic dataset that we compiled previously^59^. This dataset contains the protein sequences of phylogenetically diverse eukaryotes, and model organisms. For this project, we modified the dataset as described before^60^. An overview of the included species can be found in Supplementary Table 1, where we also report the proteome completeness scores derived from BUSCO^61^. While we attempted to remove redundancy from protein sets, some sets still contain some highly similar sequences, very likely representing isoforms encoded by the same gene. Therefore, some apparent species-specific paralogs in the identified orthologs for a given protein might not correspond to different loci, but to a single locus.

### Ortholog detection in eukaryotes

#### Protein selection and ortholog definition

As the subjects of our study, we selected the subunits of condensin I, condensin II, cohesin and SMC5/6. Additionally, we selected proteins regulating these complexes, provided that they are specific (i.e., no general cell cycle regulators like PP2A, CDK1, and Aurora kinases). The selected proteins should have been characterized as a complex component in at least one organism, predominantly *H. sapiens*, *S. cerevisiae* and *S. pombe*^28,62^, and also *T. thermophila*^24,25,63,64^*, P. patens*^65,66^*, A. thaliana*^34,67^ and *C. merolae*^68^.

For each selected protein, our objective was to detect its orthologs among the protein sequences of the species in our eukaryotic dataset. Here, ‘orthologs’ are defined as descendants from a single ancestral protein in LECA, or in a more recent lineage, if the protein is taxonomically restricted to a particular eukaryotic clade. As a result, groups of orthologs include paralogs resulting from recent duplications, that is, more recent than LECA. For example, PDS5A,B are both members of the group of PDS5 orthologs.

#### Ortholog calling: two global approaches

Each protein required its own, tailored approach to ortholog calling, of which the details can be found in Supplementary Table 2 (sheet ‘Orthologs’).

Our approaches can be globally described by two different strategies. First, for the SMC proteins SMC1-6, we searched for homologs in our eukaryotic dataset with BLASTP^69^ (E-value cutoff 1e-5) using previously identified SMC protein sequences^10^. We combined SMC protein sequences from^10^ with our hits and aligned all of them with MAFFT (v7, settings: –auto)^70^, trimmed the alignment with BMGE (v1.12, settings: -m BLOSUM30, -b 3 -h 0.7 -g 0.7)^71^ and inferred an initial a maximum likelihood phylogeny using IQ-TREE (v2.0.3, settings: -m LG+F+R5 -alrt 1000)^35^. From this phylogeny, we extracted the sequence belonging to each of SMC1-6, for which we subsequently made protein-specific profile HMMs. We used these HMMs to locally run hmmsearch (hmmer.org) across our eukaryotic dataset. We gathered the hit sequences using a domain bitscore cut-off that was well below the lowest-scoring members of putative orthologs in the initial phylogeny, to ensure all orthologs were included. For these hit sequences, we inferred a final phylogeny from which the orthologs were extracted.

Second, for non-SMC proteins, we derived profile HMMs from PANTHER17.0^72^ and EggNOG v6.0^73^. We used these HMMs to search across our eukaryotic dataset, and verified that characterized orthologs and/or functional analogs were hit by it. Then, we performed at least one iteration of the following steps: construct a profile HMM of the hits and search across our eukaryotic dataset, verify the new hits manually, and gather the trusted hits. In the next iteration, the trusted hits served as input for a new profile HMM. As long as new hits were found, we continued this cycle. For highly divergent proteins, we adopted additional strategies. These involved sequence-based strategies, like BLASTP/PSI-BLAST^69,74^, online jackhmmer^75^ and online HHpred^76^, frequently using multiple query sequences. Since sequences typically are less conserved than structures, we also adopted structure-based strategies, which entail searching for similar structures using experimentally determined structures or predicted structures from the AlphaFold Protein Structure Database^77^, applying search algorithms Dali^40^ and Foldseek^78^, also often using multiple query structures. The collected homologs were aligned to infer a maximum likelihood phylogeny (details in Supplementary Table 2, sheet ‘Orthologs’), from which the orthologs were extracted. If, because of strong sequence divergence, we expected the orthologous cluster in the phylogeny to be incomplete, we used the orthologs in the cluster to build a profile HMM to again search through our dataset, infer another phylogeny and extract the orthologs, potentially expanded compared to the previous set.

Securin and Sororin, proteins comprising intrinsically disordered regions, form a particular challenge for ortholog detection. They are highly divergent proteins, yet given their disorder, no structure-based strategy would work. A detailed description of the approach followed for Securin and Sororin can be found in the Supplementary Text.

We verified the absence of the condensin II complex from the model plant *P. patens* by searching for orthologs of CAPH2, CAPG2 and CAPD3 in the recently published near telomere-to-telomere assembly of this species^54^, which was not yet in our dataset. Indeed, we could not find candidate orthologs of these condensin II proteins in the predicted proteins of this new assembly.

We confirmed our *D. melanogaster* Nse5 candidate by first searching for its homologs with BLASTP versus NR (max. 250 hits, a permissive E-value cutoff of 1). This yielded 122 hits among a wide range of Diptera, including for instance the mosquito *Aedes aegypti* (gene 5579981, E-value=3e-09). We confirmed both the *D. melanogaster* and the *A. aegypti* Nse5 candidates to be structurally similar to Nse5 proteins in human, mouse and fission yeast using Foldseek^78^ versus AFDB-Swissprot. Of note, the IntAct^79^-reported interactors of the *D. melanogaster* Nse5 candidate seem to fulfill neural functions, and include no apparent Nse6 candidate. Thus far we have failed to detect a *D. melanogaster* Nse6 homolog.

For each protein, we provide the following data files in a GitHub repository (https://github.com/jolienvanhooff/SMCevolution):

- a text file containing the ortholog identifiers;
- a FASTA file containing unaligned orthologs;
- the profile HMM used to search for orthologs;
- the FASTA file containing the (curated) multiple sequence alignment for the profile HMM;
- the FASTA file containing the multiple sequence alignment used for phylogenetic inference;
- the logfile of the phylogenetic inference;
- the annotated gene phylogeny.

### Phylogenetic and network analysis in eukaryotes and prokaryotes

#### Eukaryotic and prokaryotic dataset assembly

To study the pre-LECA origins of eukaryotic SMC complex subunits, we assembled two protein datasets: one for eukaryotes and one for prokaryotes. The eukaryotic dataset consists of a subset of the species we used to find orthologs in, which we subsampled to accelerate the analyses and increase the interpretability of the results, while still spanning eukaryotic diversity. The selected 49 species can be found in Supplementary Table 1 (column ‘PreLECASubset’). We annotated the orthologs of our proteins of interest, to facilitate the analyses and interpretation. The prokaryotic dataset was derived from GTDB (r207) ^80^. We used a subset of the assemblies in GTDB, ultimately including one assembly for each family-level clade. For each family, we selected assemblies that A), are the GTDB representative for a given species, B), have a CheckM quality score (completeness score - 5*contamination score) above 50, and C), have the best CheckM quality score of their family. This resulted in a prokaryotic dataset containing 456 archaea and 3645 bacteria.

#### Protein selection

We selected proteins that we deemed informative about the deep evolutionary origins of SMC complexes. We thereto applied the following criteria:

1. The protein was present in LECA;
2. The protein was present in an SMC complex in LECA as a permanent constituent, i.e. not as a regulatory or transient subunit;
3. The protein belongs to a family of typical SMC complex subunits, and is not likely to be derived from other cellular machineries.

Based on these criteria we included the following proteins and their families: SMC1-6 (SMC protein family), CAPH, CAPH2, Scc1, Rec8 and Nse4 (kleisin family), CAPG, CAPD2, CAPG2, CAPD3, Scc3, PDS5 and NIPBL (hawk family), Nse1 and Nse3 (kite family), and Nse5 (family unknown) and Nse6 (family unknown). As part of our effort to find the evolutionary origins of the hawk family and Nse5,6, we searched for those of MAU2 and WAPL as well, in our HMM network analysis (see below).

#### Phylogenetic analysis

We restricted the phylogenetic analysis to the SMC proteins, the kleisins and the kites, since we deemed the hawks, Nse5 and Nse6 too divergent for phylogenetic analysis. An overview of the applied phylogenetic methods for each family can be found in Supplementary Table 2 (sheet ‘Homologs’). In general, we used profile HMM of separate orthologous sets (SMC proteins) or, for less conserved proteins, a profile HMM based on their merged alignments (kleisin, kite) to first search across our eukaryotic dataset. With this first search, we confirmed that there are no obvious other eukaryotic homologs beyond the proteins making up the SMC complexes^10^. Then, we searched iteratively across our prokaryotic sequence dataset, and collected significant prokaryotic hits. We retrieved the eukaryotic protein sequences from the ortholog sets, i.e., those belonging to species of the subsampled dataset, and combined these with the prokaryotic hits. To allow for phylogenetic inference using a relatively complex substitution model (see below), which is computationally more intense, we further downsampled the SMC and kleisin sequences based on sequence identity with CD-HIT (v4.8.1, different identity cutoffs, Supplementary Table 2, sheet ‘Homologs’)^81^. For kite, such downsampling was not necessary, because for this family we limited the phylogenetic analyses to eukaryotic sequences and their readily discernible homologs in archaea, due to which the total number of sequences was lower. We chose to exclude bacterial kites, as well as archaeal kites that we were not able to recruit with HMM-based sequence searches, since these were likely to be too distinct from the eukaryotic sequences to meaningfully align them, or to model the evolution of in the phylogenetic inference.

The remaining protein sequences were aligned using MAFFT-L-INS-i or MAFFT-DASH^82^ (v7.515), or using the PROMALS3D web server^83^. We selected the alignment approach yielding the most positions after trimming, as well as the best phylogeny (discussed below). For PROMALS3D, we obtained predicted protein structures from diverse sequences from the AlphaFold Protein Structure Database^77^, and included them as input. We trimmed the resulting multiple sequence alignments (MSAs) with trimAl in the ‘gappyout’ mode (v1.4.1)^84^ and manually cut out regions not well-conserved. We cleaned the MSA from sequences that consisted of over >70% of gaps. Phylogenetic analysis was implemented through inference of a maximum likelihood (ML) phylogeny with IQ-TREE (v.2.0.3)^35^, using the site-heterogeneous LG+C60+G substitution model and assessing branch support using ultrafast bootstrap approximation (UFBOOT^36^, 1000 replicates). To test the topology of the resulting phylogenies, we inferred another phylogeny with the same model, constraining the eukaryotic sequences to be monophyletic. We compared the original tree (the ML tree or, if it had a higher likelihood, the consensus tree) to the constrained tree to assess if the constrained tree can be rejected using the approximately unbiased (AU) test provided by IQ-TREE.

For all families, we attempted to use PROMALS3D for generating an MSA, because it, in theory, provides a means to improve the alignment of divergent sequences, since the sequence alignment is guided by the alignment of (more conserved) structures. While this worked well for the SMC proteins, it did not for the kleisin and kite families, as subsequent phylogenetic inference failed to recover the individual eukaryotic orthologous groups. Sequence members of these orthologous groups were not monophyletic, while they should have been. Moreover, ultrafast bootstrap support values were low for various key bipartitions. Possibly, these poor signals are resulting from a flawed structural guide alignment, although we did not further examine the potential underlying causes. For kites, we were able to use some structural information by employing MAFFT-DASH, which recruits resolved protein structures from the PDB, based on sequence similarity. Since it does not use predicted protein structures, the number of structures used as guides is low. For kleisins, generating a good MSA turned out particularly challenging. Therefore, we applied MAFFT-L-INS-i to the individual orthologous groups (except for Scc1 and Rec8, who were combined into a single alignment, because they were interspersed in previous phylogenies), and subsequently merged them using the ‘merge’ option, also adding the prokaryotic homologs.

PROMALS3D alignment-based phylogenies were pruned to remove leaves that corresponded to input structures. Furthermore, the SMC phylogeny, in the final visualization in Figure 4, was pruned from four leaves with very long branches (>5 average substitutions per site), and with unclear phylogenetic affiliations, since these might comprise spurious sequences. The clades were annotated according to A), their membership of a eukaryotic orthologous group (eukaryotic sequences) and B, their sisterhood relationship to the eukaryotic orthologous groups (prokaryotic sequences). If their relationship to eukaryotic sequences was not clear, we labeled them as ‘unknown’ (not necessarily monophyletic), or, in the case of the SpcB-like kites, as ‘ScpB-like’. The distributions of each group across archaea are provided in Supplementary Table 3. Note that, for SMC and kleisin proteins, these distributions form an underrepresentation due to the downsampling of the sequences for the phylogenetic analysis, as described above.

To specifically explore the relationships among the condensin/cohesin kleisins (CAPH, CAPH2, Scc1 and Rec8), we inferred an additional, more restricted phylogeny. We collected prokaryotic homologs based on a profile HMM constructed with a MAFFT-L-INS-i alignment of these four proteins. Then, we aligned the eukaryotic and prokaryotic sequences using MAFFT-LINSI and ‘merge’ as described above, trimmed the alignment with trimAl ‘gappyout’, removed sequences with >70% gaps, and applied IQ-TREE (substitution model: LG+C60+G, 1000 UFBOOT replicates). Note that in the kleisin tree, four sequences with very long branches (>5 average substitutions per site) and unclear affiliations were removed.

For the phylogenies inferred for the SMC, kleisin and kite proteins, we shared on our GitHub repository (https://github.com/jolienvanhooff/SMCevolution):

- The unprocessed multiple sequence alignment
- The multiple sequence alignment used as input for phylogenetic inference
- The maximum likelihood and the consensus phylogenies
- The logfile from the maximum likelihood phylogenetic inference
- The maximum likelihood phylogeny inferred with constraint topology, i.e. with eukaryotic sequences forced to form a monophyly.
- The reporting file of the topology test.

In addition, we share the unfiltered, uncollapsed, annotated phylogenies corresponding to Figure 4A-C on the iTOL web server^85^. Hyperlinks can be found in the legend of Figure 4.

#### Constructing the profile-versus-profile HMM network

Reconstructing the evolutionary history of the hawk proteins, Nse5,6, WAPL, and MAU2 proved challenging using classical homology detection and phylogenetic inference methods due to their high sequence divergence, as noted before^5^. To address these difficulties, we employed the profile-versus-profile HMM-based network approach introduced by^5^. Briefly, we constructed a target profile database consisting of the MSAs generated here and combined this with the precomputed COG_KOG_v1.0 database (downloaded from be http://ftp.tuebingen.mpg.de/pub/ebio/protevo/toolkit/databases/hhsuite_dbs/COG_KOG.tar.gz, last accessed Jan 17, 2023). We queried our manually-curated MSAs against this target database using the HHsearch algorithm from the HH-suite package^86^ (v3.3.0) with default parameters as implemented in the MPI Bioinformatics Toolkit^86,87^. Following this initial search, all KOG/COG profiles that were hit by a query profile (E<1000) in this first step were then used to complete reciprocal searches on the same database. The resultant graph (Supplementary Table 5) was visualized using Cytoscape^88^. Here, we first imposed thresholds for the E-value (<1e-3) and alignment length (>=100), to keep only the edges that imply meaningful significant similarity between profiles. Then, Markov Chain Algorithm (MCL) clustering was performed with an inflation parameter *I* = 3 using 1/E-value as array source as implemented in the clusterMaker2 package^89^. Finally, we used Cytoscape’s Prefuse Force Directed Layout with default settings, except for iterations = 1000, default node mass = 10, and we set the ‘Force deterministic layouts’ to ‘on’ and did not partition the graph before applying the layout. We also calculated the ‘connectivity’ between clusters. Connectivity is defined as the number of observed edges going from cluster 1 to cluster 2 (E-value=<1E-3 and alignment length>=100) over the total number of possible edges (Supplementary Table 6).

#### Hawk structural alignment

Predicted 3D structures of hawks from both *A. thaliana and H. sapiens* were directly downloaded from the AlphaFold repository (alphafold.ebi.ac.uk/, last accessed 23 December 2023). Structures were manually checked in Pymol and N- or C-terminal disordered regions and/or other domains that do not pertain to the core HEAT-repeat region were removed (see for coordinates and uniprot identifiers Supplementary Table 7). The DALI webserver^40^ was used for structural comparisons amongst the hawks, using the all-against-all function (ekhidna2.biocenter.helsinki.fi/dali). Z-scores were average clustered with the pearson coefficient as a distance measure (D=1-*r;* Supplementary Table 7). A Pymol session is included in the Supplementary Data.

#### Nse5,6 alpha-solenoid structure prediction and alignment

To predict the structures of Nse5,6 orthologs, we used the AlphaFold2 software^90^ through the local ColabFold implementation^91^. We used alignments produced by MAFFT-L-INS-i of our curated sets of orthologs as input to ensure deep alignment information to produce high-quality structure predictions. Subsequently, the alpha-solenoids were isolated from the full-length structures using the pdb_tools package^92^. The start and end coordinates of the alpha-solenoids of each full-length structure were determined by aligning them to manually-curated human Nse5 or Nse6 alpha-solenoids using Foldseek^78^. All isolated Nse5 and Nse6 alpha-solenoids were then comprehensively aligned in a pairwise fashion using TMalign^93^, where a TM-score of >=0.5 was considered the threshold for significance^94^.

### Visualization

The simplified species tree (Figures 2,6), the distributions of SMC complex subunits across eukaryotes (Figure 3) and the protein family phylogenies (Figure 4) were drawn using iTOL^85^. Phylogenies were edited and iTOL input datasets were generated using ETE (v3) ^95^ and Biopython^96^. Structures for Figure 5 were visualized and edited using PyMOL^97^. Structures for Supplementary Figure 2 were visualized using UCSF Chimera^98^. The heatmap in Figure 5B was visualized using R^99^. All final figures were prepared using Adobe Illustrator.

## Abbreviations

BRCT: <u>BR</u>CA1 <u>C</u>-terminal (repeat)
FECA: <u>F</u>irst <u>e</u>ukaryotic <u>c</u>ommon <u>a</u>ncestor Hawk
<u>H</u>EAT: protein <u>a</u>ssociated <u>w</u>ith a <u>k</u>leisin
HEAT: <u>H</u>untingtin - <u>E</u>longation factor 3 - <u>A</u> subunit of protein phosphatase 2A - <u>T</u>OR (repeat)
HGT: <u>H</u>orizontal <u>G</u>ene <u>T</u>ransfer
Kite: <u>K</u>leisin interacting tandem winged-helix <u>e</u>lement
LECA: <u>L</u>ast <u>e</u>ukaryotic <u>c</u>ommon <u>a</u>ncestor
SMC: <u>S</u>tructural <u>M</u>aintenance of <u>C</u>hromosomes
SUMO: <u>S</u>mall <u>U</u>biquitin-like <u>Mo</u>difier
TACK: <u>T</u>haumarchaeota, <u>A</u>igarchaeota, <u>C</u>renarchaeota and <u>K</u>orarchaeota
TAD: <u>T</u>opologically <u>a</u>ssociated <u>d</u>omain

## Acknowledgements

We kindly thank the following colleagues:

- Miguel Andrade-Navarro (Johannes-Gutenberg University), for his helpful advice and expert assessment on the classification of the alpha solenoid folds of Nse5 and Nse6;
- Jason Stajich (University of California Riverside) for sharing the predicted protein sequences of *Gonapodya* sp. JEL0774;
- Bastiaan de Potter, Laura van Rooijen, Carlos Santana-Molina and Berend Snel (Utrecht University) for collaborating in compiling the eukaryotic proteome dataset;
- Benjamin Rowland (Netherlands Cancer Institute), Chris Lane (University of Rhode Island) and Michelle Leger (Okinawa Institute of Science and Technology) for their constructive feedback on an earlier version of this manuscript.

This work was supported by the European Research Council (ERC Starting Grant 2018, grant number 803151, to L.E.) and the Dutch Research Council (NWO, VI.Veni.212.099 to J.J.E.H. and VI.Veni.202.223 to E.C.T.).

## Author contributions

Conceptualization, J.J.E.H., E.C.T., and L.E.; data curation, J.J.E.H., M.W.D.R., and E.C.T.; analysis, J.J.E.H, M.W.D.R., and E.C.T.; methodology, J.J.E.H, M.W.D.R, E.C.T., and L.E.; project administration, J.J.E.H.; resources, J.J.E.H, E.C.T, L.E.; supervision, J.J.E.H, E.C.T., and L.E; validation, J.J.E.H., M.W.D.R., and E.C.T.; visualization, J.J.E.H., M.W.D.R., and E.C.T.; writing – original draft, J.J.E.H. and L.E.; writing – reviewing and editing; J.J.E.H, M.W.D.R, E.C.T., and L.E.

## Supplemental Information

Document S1. Supplementary Text, Supplementary Figures 1-2, and information on Supplementary Tables 1-7 and Supplementary Data.

