## Supplementary material for "Repeated duplications and losses shaped SMC complex evolution from archaeal ancestors to modern eukaryotes": Document S1

### Supplementary Text

#### Reclassification of WAPL as ARM repeat

During eukaryogenesis, the SMC complexes were innovated with, besides hawks, other highly divergent alpha-solenoid repeat proteins, such as WAPL, MAU2 and Nse5,6. The cohesin release factor WAPL and cohesin loading factor MAU2 have been described to contain an alpha-solenoid, consisting of HEAT repeats[S1,2] and TPR repeats[S3], respectively. Whereas MAU2 was firmly affiliated with TPR repeats, strikingly, our profile similarity network revealed WAPL to be more closely related to ARM repeat proteins, instead of HEAT repeat proteins (Figure 5A). ARM and HEAT repeats, while homologous and difficult to distinguish, represent distinct evolutionary families[S4,5]. A key structural distinction is that ARM repeat units typically consist of three helices, whereas HEAT repeat units typically contain two[S4,5]. Noticeably, published structures of WAPL orthologues from human (PDB: 4K6J) and the fungus *Ashbya gossypii* (PDB: 3ZIK) typically show three helices per unit[S1,2]. When we used these structures to query the human AFDB-PROTEOME database with Foldseek[S6], we identified a strong enrichment of ARM-containing structures among the best hits (Supplementary Table 4). Consequently, we propose the alpha-solenoid present in WAPL to belong to the ARM repeat protein family, and thereby challenge the notion of a relatively recent, shared origin of WAPL with the HEAT-containing hawk proteins.

#### A shared origin of Nse5 and Nse6

The SMC5/6 complex, in addition to the SMC, kleisin and kite proteins, also contains Nse2, Nse5 and Nse6[S7]. Nse5 and Nse6 form a dimer and both predominantly feature a highly-divergent alpha-solenoid domain. Currently, there are two outstanding questions on the evolution of these proteins: firstly, whether the alpha-solenoid domains of Nse5 and Nse6 are homologous, as the similar structures of human Nse5 and Nse6 suggest, but not in those of budding yeast[S8]; and secondly, to which specific tandem repeat family these alpha-solenoid domains belong. Since Nse5 and Nse6 do not display any significant sequence similarity, we predicted their 3D structures and aligned them[S8]. Our comprehensive structural alignment analysis revealed an on-average significant similarity between predicted Nse5 and Nse6 structures from a broad sample of eukaryotic lineages (TM-score: 0.51, Supplementary Figure 2A,B)[S9], with the best alignment being highly significant (TM-score: 0.68, Supplementary Figure 2C). Altogether, this suggests that Nse5 and Nse6 are likely homologous. In the absence of any clear other homology links, we thus concluded that Nse5 and Nse6 are closest paralogs, resulting from a gene duplication before LECA.

The Nse5,6 alpha-solenoid has been hypothesized to adopt a HEAT repeat-like fold, implying homology to hawk proteins, but this classification has remained tenuous due to their high level of sequence and structural divergence[S10,11]. By leveraging our Nse5,6 orthologous sequences and predicted structures, we surprisingly found that the N-terminal repeats of the Nse5 alpha-solenoid show homology to TPR/PPR, rather than HEAT. Strikingly, the remainder of the Nse5 and the complete Nse6 alpha-solenoids show twilight-zone (1000>E-value>0.001) hits for hawk proteins

(Supplementary Table 5), suggesting an ancient homology followed by marked subsequent divergence in the Nse5 and Nse6 alpha-solenoids. Combined, we raise the possibility that the Nse5,6 alpha-solenoids are composites of highly divergent TPR/PPR and HEAT repeats, with the latter potentially being the most closely related to the hawks. Therefore, we believe it is possible that Nse5,6 emerged through duplication of an ancestral hawk protein, followed by the addition of several TPR/PPR-like repeats, and finally a pre-LECA duplication. However, it should be noted that it is unusual to have an alpha-solenoid consisting of different tandem repeat types[S12], and the identified similarity to the hawk proteins is weak, making definitive interpretations of these results difficult. Regardless of their repeat class, Nse5 and Nse6 show tremendous divergence, also within their orthologous groups, and the underlying reason remains an open question. Possibly, their divergence is driven by their role in viral restriction[S7], which implicates them in an evolutionary arms race with viruses[S13].

### Homolog detection of highly divergent proteins with functional motifs

Cohesin regulators Securin and Sororin are predominantly composed of intrinsically disordered regions, which accounts for their low conservation across eukaryotes. Consequently, for identifying their orthologs, we have primarily relied on short protein motifs often associated with protein-protein interactions. However, the mere presence of such a motif does not necessarily designate a protein as an ortholog, as these simple motifs can arise through convergent evolution.

Securin has been described in human and yeast, and a candidate has been identified in *Arabidopsis thaliana*, called Patronus[S14]. Although previous analyses did not reveal significant sequence similarity between human and yeast Securins[S15], the latest Pfam HMM profile (PF04856) encompasses both. To gather additional homologs, we searched for protein sequences containing the following motifs: a KEN box (lysine-glutamic acid-asparagine, which is involved in APC/C-mediated Securin proteolysis), a D-box (destruction box, which is also involved in APC/C-mediated Securin proteolysis), LPE (leucine-proline-glutamic acid, which blocks the interaction between Scc1 and Separase), SIS (separase-interacting segment, which inhibits Separase's protease activity by blocking substrate access to its active site)[S16–18]. Most candidate orthologs have an easily discernible KEN box and SIS motif. However, the D-box and LPE seem less conserved, and their positions within the protein seem to vary. Using iterative searches with jackhmmer[S19], we were able to connect animal and yeast Securin to plant Patronus proteins, thereby expanding the candidate ortholog set. Nonetheless, these searches also retrieved proteins like Cyclin(A/B) and Shugoshin due to their inclusion of KEN and D-box motifs. These proteins also have their own signature domains, which we searched for to exclude false Securin ortholog candidates.

Initially, we found Sororin to be metazoan-specific, but a recent paper[S20] presented compelling evidence on the homology of the C-terminus of two candidate proteins (and others) in *A. thaliana* and *S. pombe* (Sor1), specifically. This domain consists of two predicted alpha-helices in a region rich in both negatively charged amino acids (D/E) and bulky hydrophobic residues (FWL). To extend their observations we employed a strategy of profile sequence searches, whereby iterations of jackhmmer were used with different seed sequences. In case these jackhmmer iterations would either converge or share several hits along their trajectories, we concluded they would be homologous. During these iterations, we would also allow for the inclusion of so-called twilight zone hits. Such hits do not strictly fall within the category of statistically significant similarity, but would nonetheless be included because of their compelling similarities in protein length (Sororin is relatively short) and amino acid compositions. In addition, we generated various lineage-specific profile HMMs with which we searched for highly divergent sequences using hmmsearch –max (i.e., without any heuristic filter). We required a candidate ortholog to at least be able to reciprocate known orthologs (*A. thaliana*, fission yeast, human) upon several jackhmmer iterations or through a direct hmmsearch –max search. However, given the degenerated nature of the C-terminal domain/motif, it proved difficult to achieve such reciprocity and we were left with around 20 putative Sororin orthologs (e.g., in Discoba and Amoebozoa) that were not included in the final list. To our surprise, we could also find some instances of the relevant motifs in other proteins. One such instance is the kinase Haspin. In plants, it appears that Sororin duplicated and one of the copies was fusion to the N-terminus of Haspin kinase. Such findings suggest a dynamic evolution of Sororin's signatures, which might be searched for across a wider range of proteins, as they may represent similar functions or mechanisms.

#### Genomic co-localization of archaeal SMC complex subunits separates a condensin/cohesin-related complex and an SMC5/6-related complex

We hypothesized that many archaea, in particular TACK and Asgard archaea, have two SMC complexes, both with their own SMC, kleisin and kite protein, where one complex is closer related to condensin and cohesin (while retaining a kite protein instead of a hawk protein), and the other to SMC5/6. To test this hypothesis, we explored the localization patterns of the genes encoding these proteins in archaeal genomes. After all, if these subunits form individual complexes, one would expect the genes of a single complex to co-localize in the genome, presumably in a single operon. While our sets of archaeal SMC proteins and kleisins are limited (since we downsampled these data to facilitate the phylogenetic analysis, see Methods), we still have several dozens of species in which we identified more than one subunits, so we could check whether they co-localize.

For condensin/cohesin-related subunits, we inspected the SMC proteins that form a sister clade to SMC1-4 and those of unknown identity (Figure 4A, "SMC1-4-related" and unlabelled archaeal clades), as well as the kleisin subunits of unknown identity (Figure 4B, unlabelled archaeal clades), and the ScpB-like kite proteins (Figure 4C). 22 of the 38 species having both such an SMC and a kleisin protein, have these subunits directly adjacent to one another in their genomes, and 79 out of the 124 species with both such a kleisin and kite protein have these as neighbors (Figure 4D, Supplementary Table 3). The SMC and kite are found directly next to one another only in one out of 48 species, since the kleisin gene often sits in between. Yet, we might often not have identified this kleisin due to downsampling. For example, *Prometheoarchaeum syntrophicum*, one of the only two Asgard archaea that have been cultured[S21], has a genomic region with the condensin/cohesin-related SMC, kleisin and kite next to one another, in that order (assembly NCBI RefSeq assembly GCF_008000775.1, genes NZ_CP042905.1_810, NZ_CP042905.1_809 and NZ_CP042905.1_808 as derived from GTDB). In addition to supporting their complex formation, these colocalizations also confirm that the archaeal kleisins of unknown identity, as discussed in the main text, correspond to condensin/cohesin-related kleisins, as we proposed, and that their placement in the phylogeny (Figure 4B) is incorrect, as they should be the sister-group to CAPH+CAPH2+Scc1+Rec8. Moreover, it offers the opportunity to screen for other condensin/cohesin-related SMC, kleisin or kite subunits that we thus far have not identified, by examining the proteins neighboring already identified subunits.

For SMC5/6-related subunits, we inspected the SMC proteins that form a sister clade to SMC5-6 (Figure 4A, "SMC5-6-related"), the kleisin proteins that form a sister clade to Nse4 (Figure 4B, "Nse4-related") and kite proteins that cluster with or within Nse1 and Nse3 (Figure 4C, "Nse1-3-related" and "Nse3-related"). In two out of nine species with such an SMC and kleisin protein, these genes are next to one another. Strikingly, in all 31 instances in which we identified both an SMC5/6-related kleisin and kite gene, they neighbor each other. Under our strict requirement for being direct neighbors, the SMC5/6-related SMC protein and kite protein of *P. syntrophicum* are not counted as co-localizing, since they are interspersed by one gene (NCBI RefSeq assembly GCF_008000775.1, genes NZ_CP042905.1_495 and NZ_CP042905.1_497, respectively). We inspected their in-between gene (NZ_CP042905.1_496), and it turned out an Nse4-related kleisin, which we had excluded while downsampling sequences for phylogenetic analysis. Hence, in addition to a complete co-localizing set of condensin/cohesin-related subunit genes, *P. syntrophicum* also has a complete co-localizing set of SMC5/6-related subunit genes (Figure 4D). This confirms that gene localization patterns can be used to identify additional subunits in archaea, but, more importantly, it confirms the existence of two archaeal SMC complexes. Likewise, we regard it unlikely that the archaeal SMC protein related to SMC1-4 and the one related to SMC5-6 form a heterodimer together, as discussed in our evolutionary scenario for the origins of the four eukaryotic SMC complexes.

In addition to confirming the existence of the two ancient archaeal SMC complexes, the genomic context also contains information about the intriguing option that the predecessor of Nse1 and Nse3 already duplicated in an Asgard archaeal ancestor of eukaryotes, and hence is not eukaryote-specific. Five Asgard archaea have both an Nse3-related kite and a generic Nse1-3-related kite (Figure 4C). Two out of these five Asgard archaea, both Heimdallarchaeia belonging to the clade of Hodarchaeales, have multiple SMC5/6-related SMC proteins, of which at least one SMC gene closely localizes to the Nse3-related kite, and one other to the Nse1-3-related kite. For example, Candidatus Heimdallarchaeota archaeon LC_3 (GenBank assembly GCA_001940645.1) two such SMC-kite combinations exist, but for none we had identified potential Nse4-related kleisin proteins, not even in the unsampled dataset. However, we screened the in-between genes of both SMC-kite combinations, and concluded that one of them has kleisin. Hence, this hodarchaeon probably has two sets of co-localizing SMC5/6-related subunits, one of which has a kite that seemed specifically related to Nse3 (MDVS01000027.1_69, MDVS01000027.1_68 and MDVS01000027.1_67, corresponding to the SMC, kleisin and kite gene, respectively) and another that another belonging to the large group of kites that are similarly related to Nse1 and Nse3 (MDVS01000144.1_29, and MDVS01000144.1_20 for the SMC and kite protein, respectively). While we cannot be certain about the specific relationship of these SMC5/6-related kite proteins to their eukaryotic relatives Nse1 and Nse3 (main text, "Eukaryotes inherited an expanded SMC array from their archaeal ancestors"), these observations suggest that an Asgard archaeal ancestor underwent a complete duplication of the SMC5/6-related complex, resulting in a total of three SMC complexes in this lineage. Moreover, it suggests that the kite proteins resulting from this duplication, unlike Nse1 and Nse3, are involved in different SMC complexes, whereas Nse1 and Nse3 both join the (single) eukaryotic SMC5/6 complex.

### Supplementary Figures

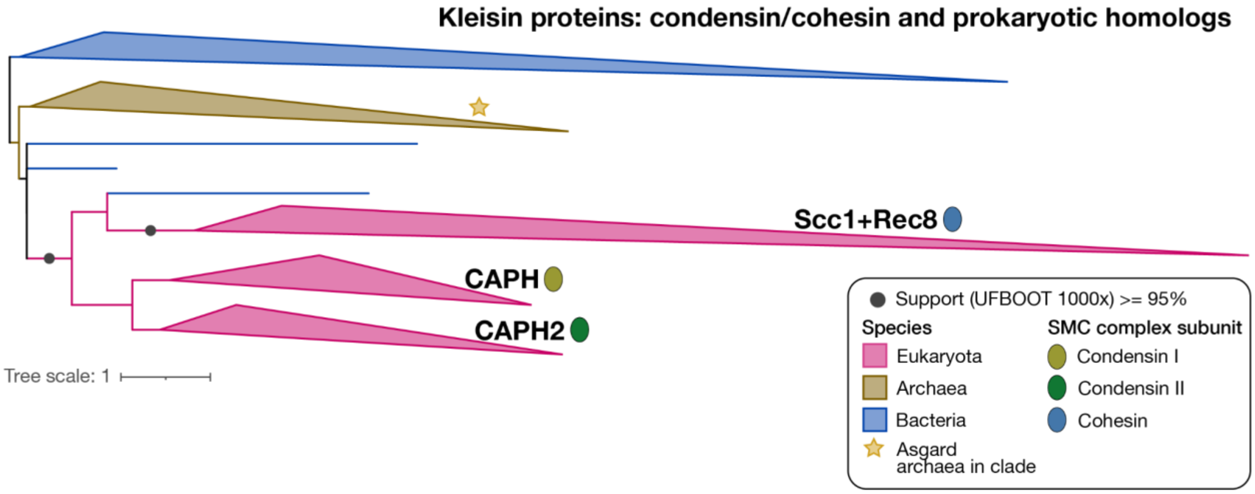

**Supplementary Figure 1. Condensin/cohesin kleisin phylogeny including prokaryotic homologs**. Inferred with IQ-TREE[S22] using the LG+C60+G mixture model. Related to Figure 4B. For the uncollapsed tree, see https://itol.embl.de/tree/892051281581924916986660 65.

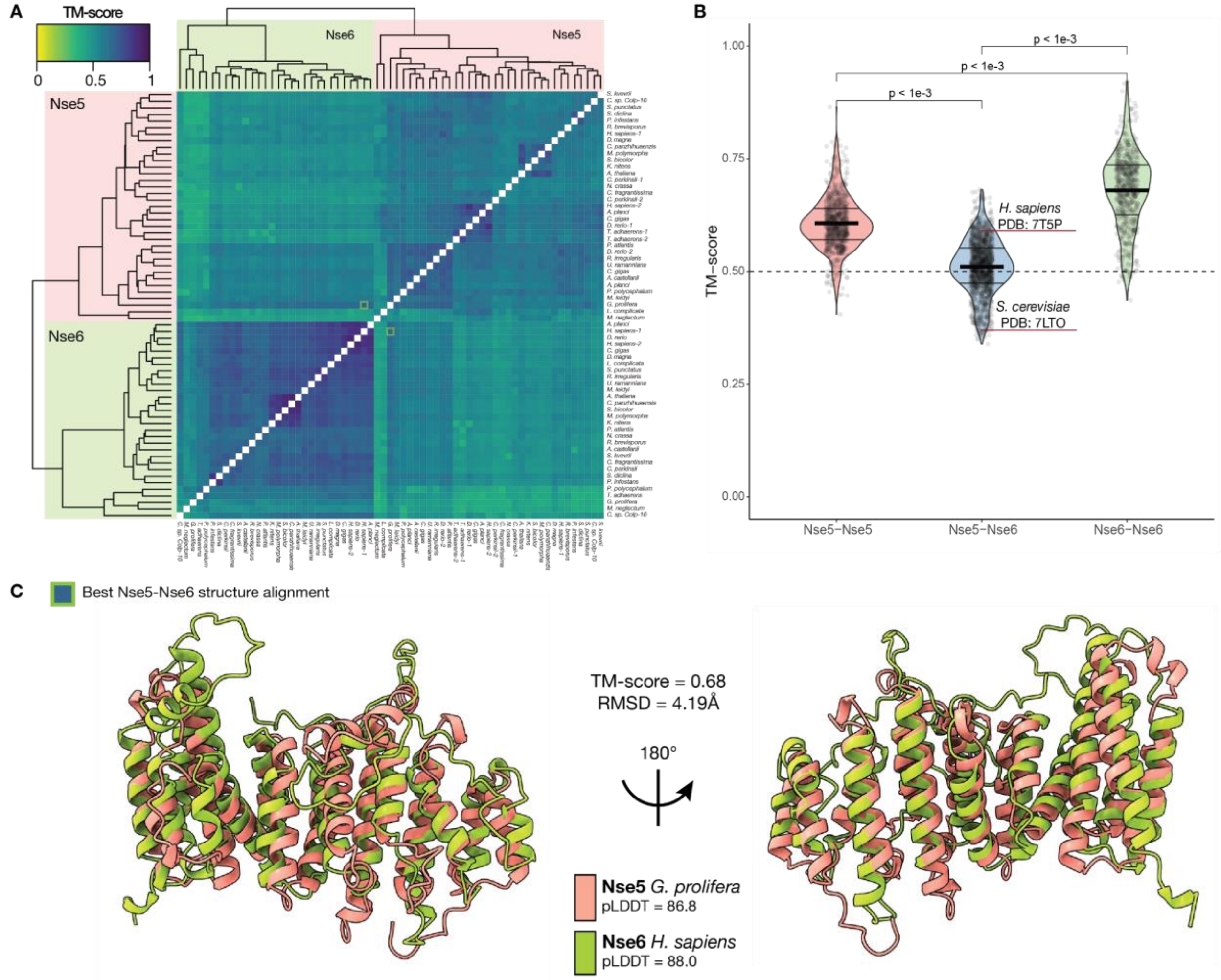

**Supplementary Figure 2. Pairwise alignment scores (TM-scores) of Nse5 and Nse6 predicted alpha-solenoid structures for species having both proteins**. **A**. Heatmap displaying the pairwise TM-scores from the structure alignment generated by TM-align[S23]. Structures are clustered based on the similarities of their TM-scores. **B**. Violin plots of TM-scores in (A) among Nse5, between Nse5 and Nse6, and among Nse6 predicted structures. Highlighted are the scores from Nse5-Nse6 structure alignments of the human[S8] and yeast (cryo-)EM structures[S11], respectively, along with their PDB entries. P-values were obtained from Student's t-tests. The dashed line marks the TM-score significance cut-off of 0.5[S9]. **C**. Structure alignment of the predicted structures of *Gonapodya prolifera* (Nse5) and *Homo sapiens* (Nse6), which presents the best alignment across all pairwise alignments in A. In addition to the TM-score, the root mean square deviation (RMSD) is shown. The predicted local distance difference test (pLDDT) indicates the quality of the structure predictions generated by AlphaFold[S24].

### Supplementary Tables

**Supplementary Table 1**. Eukaryotic taxa included in our analyses and numbers of orthologs for each of the proteins studied in their predicted proteomes.

**Supplementary Table 2**. Overview of applied methods to find orthologs of the studied proteins, and the more ancient homologs dating back before LECA, for a subset of protein families.

**Supplementary Table 3**. Presences and absences of SMC complex subunits in archaea, and the example SMC complex gene clusters in Asgard and TACK archaea (Figure 4D), as well as two detected in Euryarchaeota.

**Supplementary Table 4**. Overview of structure hits of WAPL proteins (human, *Ashbya gossypii*) in search against human predicted structures.

| **Query: 4K6J_A - Human**[S1] | | | | | |
|---|---|---|---|---|---|
| **Hit name** | **Hit ID** | **E-value** | **Probability** | **Orthologous group** | **Repeat type + reference** |
| Beta-catenin-like protein 1 | Q8WYA6 | 2.48E-04 | 1 | KOG2734 | Armadillo[S25] |
| Synembryn-B | Q9NVN3 | 1.21E-02 | 1 | KOG4464 | Armadillo[S26] |
| Plakophilin-2 | Q99959 | 2.33E-03 | 1 | KOG1048 | Armadillo[S27] |
| Armadillo repeat-containing protein 6 | Q6NXE6 | 1.31E-02 | 1 | KOG4199 | Armadillo[S28] |
| Protein SAAL1 | Q96ER3 | 1.31E-02 | 1 | ENOG502QS5W | Armadillo[S29] |
| Transmembrane and coiled-coil domain-containing protein 6 | Q96DC7 | 2.73E-03 | 1 | ENOG502QPMC | Armadillo[S30] |
| Importin-11 | Q9UI26 | 9.97E-03 | 1 | KOG1993 | HEAT[S31] |
| HEAT repeat-containing protein 6 | Q6AI08 | 6.47E-03 | 1 | KOG4535 | HEAT[S31] |
| Catenin delta-2 | Q9UQB3 | 5.53E-03 | 1 | KOG1048 | Armadillo[S32] |
| Telomere repeats-binding bouquet formation protein 1 | Q8NA31 | 2.27E-02 | 1 | ENOG502QQER | Armadillo[S31] |
| APC2 | O95996 | 3.79E-02 | 1 | KOG2122 | Armadillo[S33] |
| Protein zer-1 homolog | Q7Z7L7 | 1.42E-02 | 1 | KOG3665 | Armadillo[S34] |
| **3ZIL_A-*Ashbya gossypii***[S2] | | | | | |

| Hit output | Hit ID | E-value | Probability | Orthologous group | Repeat type |
|---|---|---|---|---|---|
| Telomere repeats-binding bouquet formation protein 1 | Q8NA31 | 1.73E-01 | 0.99 | ENOG502QQER | Armadillo[S31] |
| Tetratricopeptide repeat protein 12 | Q9H892 | 1.06E-01 | 0.99 | KOG0548 | TPR/Armadillo |
| Protein zyg-11 homolog B | Q9C0D3 | 2.62E-01 | 0.99 | KOG3665 | Armadillo[S35] |
| Plakophilin-1 | Q13835 | 9.74E-02 | 0.99 | KOG1048 | Armadillo[S27] |
| Beta-catenin-like protein 1 | Q8WYA6 | 2.13E-01 | 0.98 | KOG2734 | Armadillo[S25] |
| Catenin beta-1 | P35222 | 2.22E-01 | 0.97 | KOG4203 | Armadillo[S36] |
| Armadillo repeat-containing protein 2 | Q8NEN0 | 3.22E-01 | 0.97 | KOG1048 | Armadillo[S37] |
| E3 ubiquitin-protein ligase TRIP12 | Q14669 | 2.96E-01 | 0.97 | KOG0168 | Armadillo[S38] |
| Armadillo repeat-containing protein 8 | Q8IUR7 | 3.09E-01 | 0.96 | KOG1293 | Armadillo[S39] |
| Junction plakoglobin | P14923 | 5.06E-01 | 0.93 | KOG4203 | Armadillo[S34] |
| Brefeldin A-inhibited guanine nucleotide-exchange protein 1 | Q9Y6D6 | 1.73E-01 | 0.93 | KOG0929 | Armadillo[S40] |
| Telomere-associated protein RIF1 | Q5UIP0 | 2.13E-01 | 0.92 | ENOG502QV6C | Proposed mix armadillo/heat[S41] |

**Supplementary Table 5**. Combined results of the HHsearch runs, with probability, E-value, bitscore, alignment length, and hit type, defined as either ‘best hit’, ‘second hit’, up until ‘tenth hit’, and otherwise the generic term ‘hit’. Finally, the combined GO terms associated with the human orthologs of each KOG are listed.

**Supplementary Table 6.** Pairwise ‘connectivity’ between all pairs of MCL clusters. Connectivity is defined as the number of observed edges going from cluster 1 to cluster 2 (E-value =< 1e-3 and alignment length >= 100) over the total number of possible edges. The orthologous groups that comprise the clusters in a given pair are stated in cluster1_IDs and cluster2_IDs, respectively.

**Supplementary Table 7.** DALI Z-scores for comparisons of the 3D predicted structures of hawks of *A. thaliana* and *H. sapiens*. Links to the AlphaFold2 predicted structures are embedded, including their average pLDDT score, UniProt ID and the coordinates of the domains that were used for the structural comparison.

### Supplementary Data

Data related to the orthologs of the SMC complex subunits, to the origins of the eukaryotic SMC complexes, and the metadata on our eukaryotic and prokaryotic datasets can be found in a GitHub repository (https://github.com/jolienvanhooff/SMCevolution, see README.md for a description of the data). Note that the ortholog sets of the subunits may contain some redundancy, specifically seemingly species-specific paralogs corresponding to isoforms (Methods, ‘Eukaryotic dataset assembly’). Uncollapsed phylogenies corresponding to Figure 4A-C can be found on iTOL (5A, SMC proteins: https://itol.embl.de/tree/13722425212266531697468040; 5B, kleisin family: https://itol.embl.de/tree/137224252126461697465203, 5C, kite family: https://itol.embl.de/tree/137224252121987116975 25255).

### Supplementary Abbreviations

| | |
|---|---|
| APC/C | Anaphase-Promoting Complex/Cyclosome |
| ARM | Armadillo (repeat) |
| BRCT | BRCA1 C-terminal (repeat) |
| Hawk | HEAT protein associated with a kleisin |
| HEAT | Huntingtin - Elongation factor 3 - A subunit of protein phosphatase 2A - TOR (repeat) |
| Kite | Kleisin interacting tandem winged-helix element |
| LECA | Last eukaryotic common ancestor |
| PPR | Pentatricopeptide repeat |
| SMC | Structural Maintenance of Chromosomes |
| TACK | Thaumarchaeota, Aigarchaeota, Crenarchaeota and Korarchaeota |
| TPR | Tetratricopeptide repeat |
